# scBiopsy-seq: a platform for temporal single-cell RNA-seq analysis

**DOI:** 10.1101/2025.03.26.645409

**Authors:** Linfeng Cai, Shiyan Lin, Minghao Qiu, Li Lin, Fuyuan Li, Jiajia Liu, Yuning Zou, Xing Na, Shanshan Liang, Xing Xu, Chaoyong Yang, Jin Li

## Abstract

The development of temporal single-cell RNA-seq (scRNA-seq) assay enabled us to systemically investigate the effects of various types of perturbations in a time course with single-cell resolution. However, the existing temporal scRNA-seq technologies have certain limitations in reliability, detection efficacy and detection diversity. In the current study, we develop scBiopsy-seq assay, which combines a synergistic electroosmosis-electrophoresis extraction method for efficient RNA extraction and digital microfluidics with contamination-isolated hydrophobic interface for high-performance sample processing. scBiopsy-seq extracts the cytoplasm at well-controlled volume to detect >10K genes per extraction with 90% successful rate. Functional enrichment analysis revealed that the genes robustly detected by scBiopsy-seq were associated to diverse biological processes, demonstrating its superb diversity of detection. scBiopsy-seq can perform the sequential extraction of the cytoplasm from the same cell multiple times, which allows us to associate the cell phenotypic responses with its transcriptional dynamics. We employed scBiopsy-seq to analyze the temporal response to a BRD4 degrader induced transcriptional suppression, which identified the key roles of fatty acid beta oxidation in this biological process. scBiopsy-seq with similar data quality as scRNA-seq will dramatically expand the application of temporal scRNA-seq towards a broader spectrum of cell biology research.

## Introduction

The single-cell technology to measure various types of signals including transcriptome, methylome or epigenome with single-cell resolution has been developed rapidly in the past decades. Many efforts have been spent to establish assays to perform two, three or even more omics analysis within the same cell. These assays can measure transcriptome-methylome, transcriptome-methylome-epigenome or non-sequencing derived data with single-cell resolution ^1-6^. These efforts have been expanded to the two-dimension world, measuring transcriptome alone or transcriptome-epigenome spatially with single-cell resolution ^7-9^. In addition, experimental or computational methods have also been built to integrate the information of mitochondrial transcriptome into the regular single-cell RNA sequencing (scRNA-seq) analysis for broader applications ^10,11^. In comparison, more efforts are still needed for the development of temporal scRNA-seq technologies. It is important to understand the diversity of a group of cells, but it is even more important to understand how this diversity can impact the cellular functions. To analyze the changes of cell biology responding to stimulations within the same cell in a time course is the key to understand the functional significance of intercellular diversity.

The differences of responses to stimulation among cells are decided by the features of cells before the stimulation was applied. Upon stimulation, the features of responding dynamics in every cell demonstrate the diversity of responses. Currently, many scientists rely on their biological expertise to identify and validate the potential important features based on the snapshots of the single-cell transcriptome at each time point. The establishment of computational algorithm such as pseudotime ^12^ or RNA velocity ^13^ helps us to identify the association of different cells with single-cell resolution in a time course upon stimulation. However, the trajectories as well as the underlying features discovered by the computational algorithms are rather hints or suggestions, which require intensive experimental validation. The metabolic labeling based method may distinguish the newly-generated mRNAs from old mRNAs in single-cell resolution ^14^, but currently it cannot analyze the transcriptome of the same cell in a time course. Therefore, a temporal scRNA-seq technology, which can provide the information of the transcriptome at each time point from the same cell, is desired for universally biological applications.

A few methods ^15,16^, including Live-seq, have been established to profile the changes of transcriptome in a time course within the same cell. Despite of the revolutionary contribution, the applications on biological research have been limited due to various kinds of reasons. The mechanic-driven extraction method leads to the uncontrollable extraction volume over a wide range, which may be a disturbing factor contributing to its undesirable success rate (40%). As the average number of genes detected by each extraction is ∼2100 at the sequencing depth of 1 million, remarkable drop-out rate for the data from Live-seq is expected. In the current study, we established an assay, so called scBiopsy-seq, to achieve well-controlled extraction volume, high detection efficacy and high reliability for the measurement of transcriptome for the same cell in a time course.

## Results

### The designing of scBiopsy-seq

We aimed to design a platform, named as scBiopsy-seq, to measure the transcriptome at various time points of the same cell. The goal of scBiopsy-seq is to achieve the similar reliability and efficacy as the regular Smart-Seq2 (SS2) based scRNA-seq for every extraction. We proposed to use nanopipettes driven by electronic field, based on a three-electrode system, to achieve accurate control of extraction volume for enhanced stability and successful rate in temporal scRNA-seq. A nanopipette with ∼ 300 nm needle tip was used for extraction to minimize cellular perturbation (**Supplementary Fig. 1A**). The methods for creating these nanopipettes were provided in **Supplementary Methods and Supplementary Data Fig. 1B**.

It has been described that the electroosmosis (EOF) flow generated by the negatively charged silanol surfaces of the nanopipettes towards the cathode can extract intracellular components from cells ^17^. However, the electrophoretic force (EP) acting on negatively charged RNA molecules towards the anode, which is in the opposite direction of the EOF, may impair the extraction efficacy ^18^. We hypothesized that this issue can be solved by adding the 3-Aminopropyltriethoxysilane (APTES) modification on the nanopipettes to form positively charged surfaces, which may convert the direct the EOF towards anode in order to generate synergistic effects of the EOF and EP (**Supplementary Fig. 2A)**.

The pipette-based liquid handling for the library construction may lead to dramatic loss of samples. In our previous studies, we have confirmed that the application of pipette-free methods, such as the digital microfluidics (DMF) assays, can significantly improve the efficacy of single-cell sequencing ^19,20^. Since the oil-isolated hydrophobic interface on the DMF can reduce the nucleic acid adsorption and avoid exogenous contaminants, we hypothesized that the integration of the DMF for the fully automatic RNA sample processing (**Supplementary Fig. 2B-D**) may further increase the efficacy and successful rate of temporal scRNA-seq.

The scBiopsy-seq assay was established based on the above two hypothesis. The cells were kept in an environmental chamber with 5% CO_2_ in 37°C and visualized by a regular laboratory microscope. For each experiment, the nanopipette was inserted into one cell and the cytoplasm extraction was performed by the synergistic EOF and EP with the three-electrode system. The samples were then transferred from the nanopipette into the microfluidics chip for a fully automatic processing ^21^. In our hands, a trained biologist can perform one experiment within 5 min with >90% successful rate (138 of 150 tests by four biologists independently), with comparable throughput as Live-seq (5 extractions per hour). The design of scBiopsy-seq assay was presented at **Figure 1A**. The instruments for scBiopsy-seq were presented at **Figure 1B**.

**Fig. 1.**
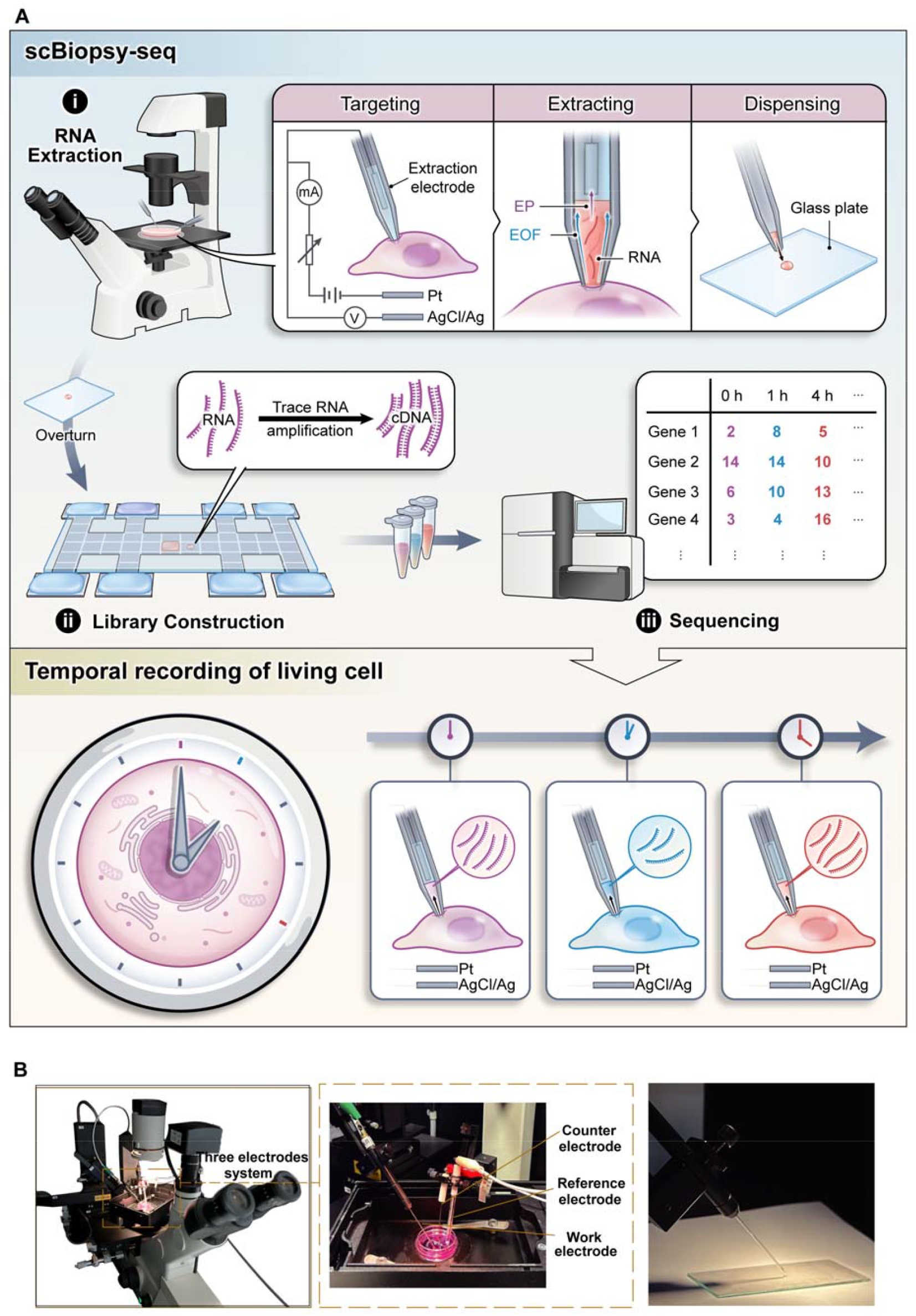
The design of scBiopsy-seq for temporal single-cell RNA-seq. **A**. The working principle of scBiopsy-seq. scBiopsy-seq assay was based on the microinjection-extraction instrument for RNA extraction and digital microfluidics for RNA library processing; **B**. The three-electrode assay for RNA extraction and transfer.

### scBiopsy-seq extracts cell cytoplasm from individual cell with high efficacy and precise volume control

The hypothesis that aligning EOF and EP to the same direction may increase the extraction efficacy was tested virtually by the simulation for fluid physics with COMSOL Multiphysics (https://www.comsol.com/comsol-multiphysics) (**Supplementary Methods**). In comparison to the results using nanopipette with negatively charged surface (**Fig. 2A**), the APTES-modified positively charged surface generating the synergistic EOF and EP can improve the extraction efficiency, demonstrated by the elevated concentration field of negatively charged substance including RNAs (**Fig. 2B-C**). Furthermore, with the increase of positive charge from 70 mC/m^2^ to 240 mC/m^2^ on the surface, the average concentration of negatively charged substance within the nanopipette increases from 367.52 mM to 1215.40 mM (**Supplementary Fig. 2E-G**). These results suggested that the synergistic EOF and EP in the same direction may be an important factor for temporal scRNA-seq analysis.

**Fig. 2.**
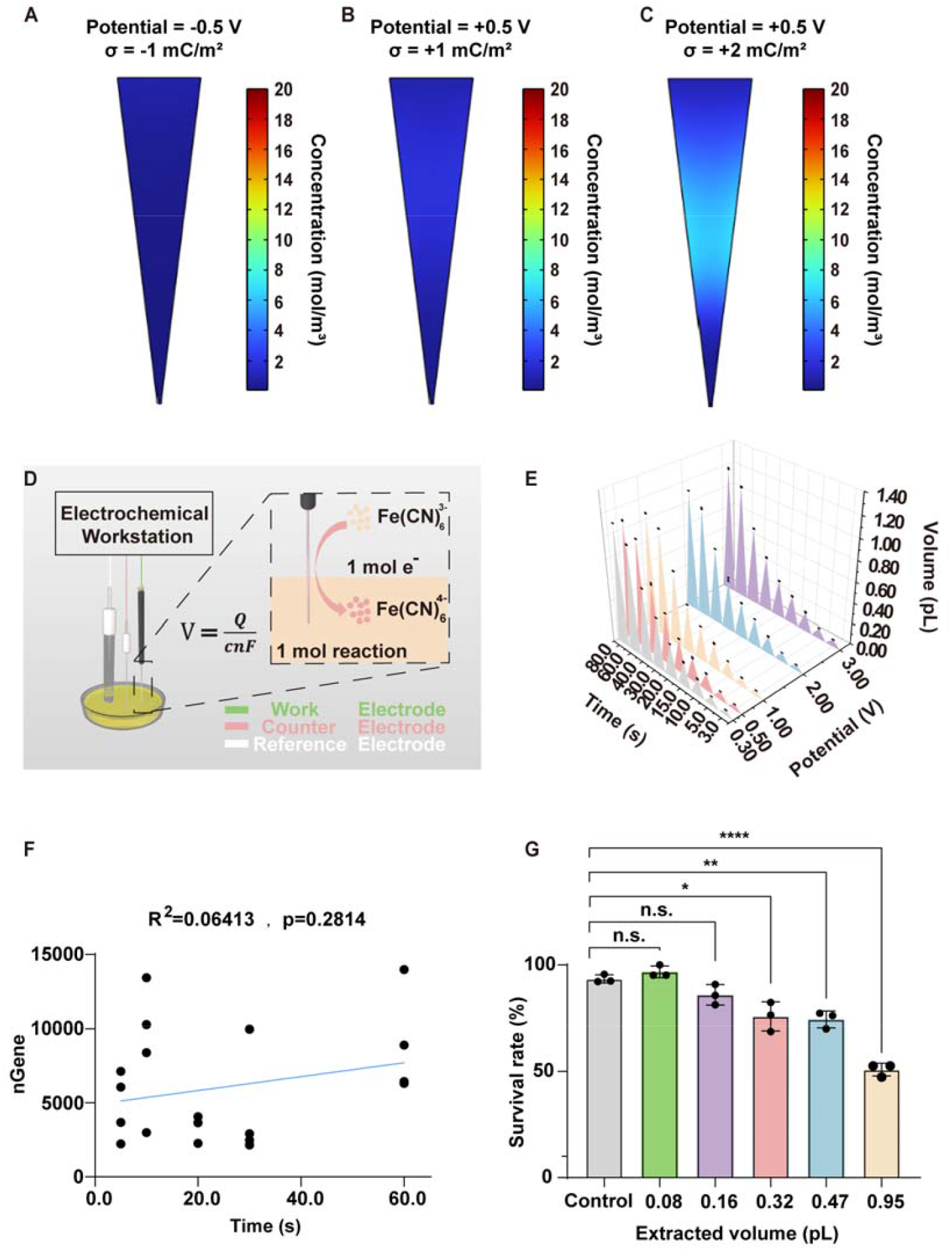
scBiopsy-seq can extract samples with well-controlled volume. **A**. Fluid physics simulation of concentration field with COMSOL Multiphysics in a capillary during the extraction process. The charge surface density (σ) was –1 mC/m2 and the corresponding applied potential was –0.5 V. The extracting concentration of negatively charged molecules, such as RNAs, was presented in color scale. Average concentration of negatively charged substance within the nanopipette was calculated at 0.51 mM. **B** Fluid physics simulation of concentration field with COMSOL Multiphysics in a capillary during the extraction process. The σ was 1 mC/m2 and the corresponding applied potential was +0.5 V. The extracting concentration of negatively charged molecules, such as RNAs, was presented in color scale. Average concentration of negatively charged substance within the nanopipette was calculated at 1.49 mM. **C**. Fluid physics simulation of concentration field with COMSOL Multiphysics in a capillary during the extraction process. The σ was 2 mC/m2 and the corresponding applied potential was +0.5 V. The extracting concentration of negatively charged molecules, such as RNAs, was presented in color scale. Average concentration of negatively charged substance within the nanopipette was calculated at 3.75 mM. **D**. Schematic of the experimental design to assess extraction volume. When 1 mol of potassium ferrocyanide is involved in the reaction, the charge transferred at the electrode amounts to 1 mol, enabling the calculation of extraction volume according to Faraday’s law. **E**. The extraction volume determined by different applied voltages and extraction time. N=5. **F**. The correlations (R^2^ of linear regression and p value (two-sided F-test)) between the number of detected genes and extraction time. **G**. The survival rate of SK-BR-3 cells under different extracted volumes. Three groups were tested and 20 cells were contained in each group.

As we used EOF and EP for RNA extraction, it is possible to control the extraction volume precisely in comparison to the mechanic-driven assays such as Live-seq. This hypothesis was tested with the Fe(CN) ^4-^ solution as the mimic for cytoplasm according to the Faraday’s Law ^22^ (**Fig. 2D**). The results indicated a controllable volume (0.05 pL-1.13 pL) determined by extraction voltage and time with CV<4.08% (**Supplementary Methods and Fig. 2E**). Especially, we observed a linear correlation between extraction volume and time at a given voltage (**Supplementary Fig. 3A**), which enables us to perform the extraction at precise volume.

Then we investigated the effects of well-controlled extraction volume on the measurement of transcriptome in SK-BR-3 cells. The variation of extraction time may have negligible effects on the number of detected genes and counts (**Fig. 2F and Supplementary Fig. 3B**). Furthermore, the extraction volume had no obvious effects on the clustering of transcriptome data (**Supplementary Fig. 3C and Supplementary Table 1**). Then we tested if the extraction volume had any effects on the viability of SK-BR-3 cells. Indeed, we observed the decreases of survival rate when the extraction volume was larger than 0.16 pL (**Fig. 2G**), which accounted for ∼10% of the average volume of SK-BR-3 cells (**Supplementary Fig. 3D**). These results indicated that some of the extractions with volume >0.16 pL, which is likely to happen when the extraction volume cannot be controlled, may hurt the cell viability and cause the failure of the entire experiments. Actually, the successful rate of scBiopsy-seq by a trained biologist is >90% (138 of 150 tests by four biologists independently), which is much higher than that of Live-seq (40%). Overall, scBiopsy-seq can extract samples from individual cell with high efficacy and reliability.

### scBiopsy-seq performs single-cell temporal transcriptome analysis with high gene detection capacity and functional diversity

Based on the encouraging results of benchmarking for the extraction method, we continued to benchmark the performance of scBiopsy-seq in measuring the transcriptome. We firstly confirmed that the presence of current flow derived by the electronic force was vital for RNA extraction, as we can only detect 20-40 genes per extraction without current flow (**Supplementary Fig. 4A**). With the application of synergistic EOF-EP as well as microfluidics-based library construction, scBiopsy-seq can detect ∼10K genes per extraction reproducibly, which is significantly improved in comparison to Live-seq (>3-fold) or scBiopsy-seq without synergistic EOF-EP (>2-fold) (**Figure 3A and Supplementary Table 2**). In detail, scBiopsy-seq in average can obtain the information of 60.3% uniquely mapped reads, 211399 counts and 10460 genes per extraction on average with a low percentage of the reads from mitochondrial genome (**Figure 3B and Supplementary Table 3**). The covalent modification of APTES on the nanopipette is essential, as the number of detected genes decreased dramatically when APTES was added into the lysis buffer directly (**Supplementary Fig. 4B and Supplementary Table 4**).

**Fig. 3.**
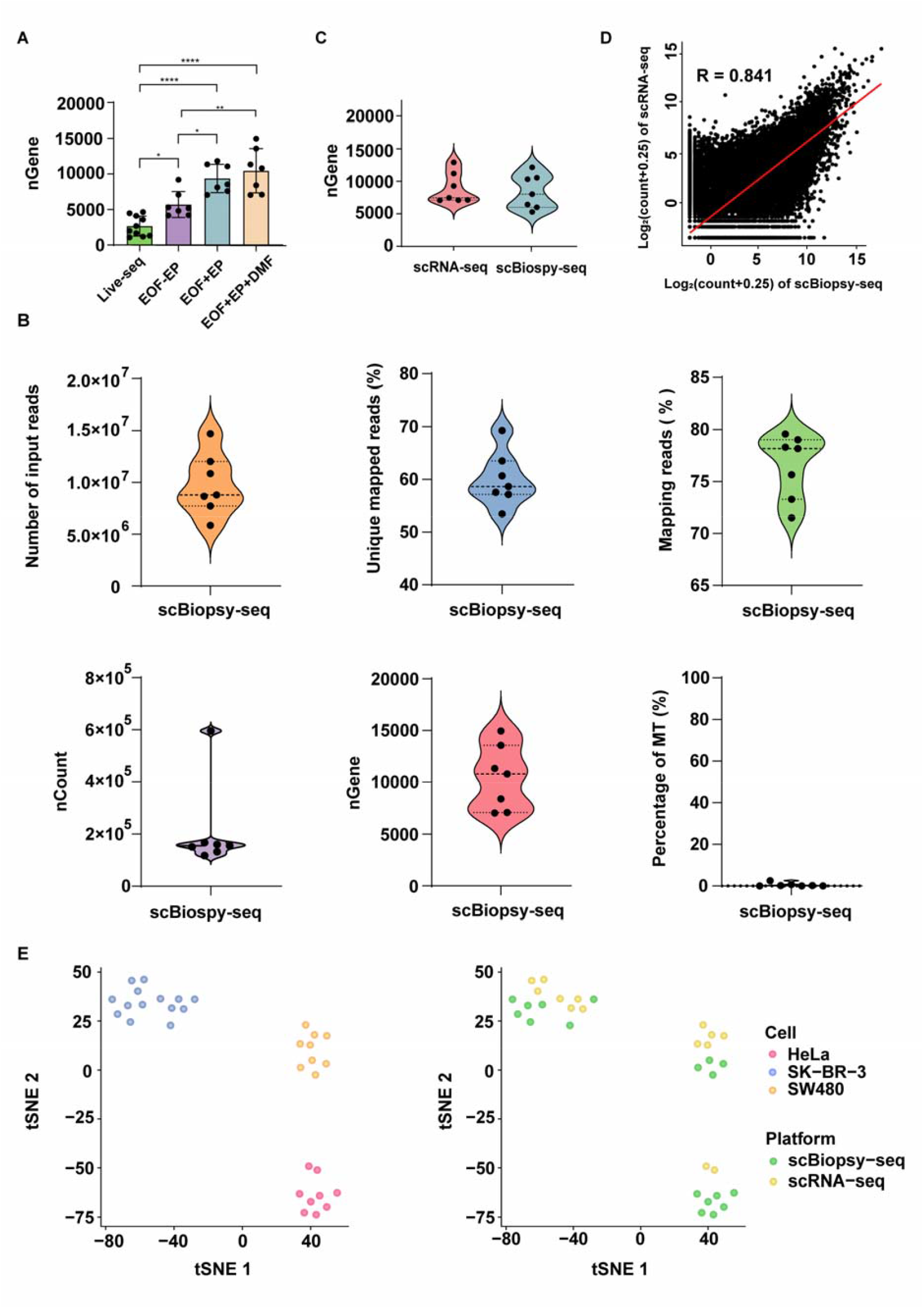
scBiopsy-seq can measure the transcriptome reliably and efficiently. **A**. The number of genes detected in single RAW264.7 cells by using different methods and conditions. All the data were downsampled at the sequencing depth of 1 million reads for fair comparison. N=10. EOF-EP: no APTES was modified on the pipette; EOF+EP: APTES was covalently modified on the pipette; EOF+EP+DMF: APTES was covalently modified on the pipette and the RNA library was processed by DMF device. **B**. Basic parameters of scBiopsy-seq applied on RAW264.7 cells. N=7. nGene: the number of detected genes; Percentage of MT: percentage of reads from mitochondrial genes. Counts and nGenes were calculated at the sequencing depth of 1 million reads. **C**. The number of genes detected in SK-BR-3 cells by Smart-seq2 based scRNA-seq and scBiopsy-seq. N=7. **D**. The correlation between the transcriptome data obtained from scBiopsy-seq assay and Smart-Seq2 based scRNA-seq assay. N=7. **E**. tSNE map of data from scRNA-seq or scBiopsy-seq from three cell lines by unsupervised clustering. The results were presented based on different cell lines (**left**) and different assays (**right**).

We then compared the detection efficacy of scBiopsy-seq and regular scRNA-seq. The numbers of detected genes were comparable between scBiopsy-seq and SS2 based scRNA-seq (**Fig. 3C and Supplementary Table 5**). The expression of genes across the whole transcriptome, from scBiopsy-seq assay or scRNA-seq, presented a significant correlation (**Fig. 3D**). The transcriptome data of three different cell lines, from either scBiopsy-seq or scRNA-seq, were clustered properly (**Fig. 3E and Supplementary Table 6**). These results revealed that scBiopsy-seq can obtain the information of transcriptome with excellent detection efficacy from single extraction, which is comparable to the results of SS2 based scRNA-seq from a whole cell.

The fidelity of scBiopsy-seq was validated with multiple assays. The detected genes of scBiopsy-seq were distributed evenly in different chromosomes without dramatic bias (**Supplementary Fig. 4C**). A general concern for temporal scRNA-seq assays is whether the extraction operation itself has dramatically change the transcriptome, which may impair the fidelity of the analysis in a time course. We therefore investigated the transcriptome from multiple extractions of SK-BR-3 cells. Encouragingly, the transcriptome data from different extractions in a time course presented a tight correlation (**Fig. 4A and Supplementary Table 7**), suggesting the extraction operation did not induce the significant alterations of the transcriptome.

**Fig. 4.**
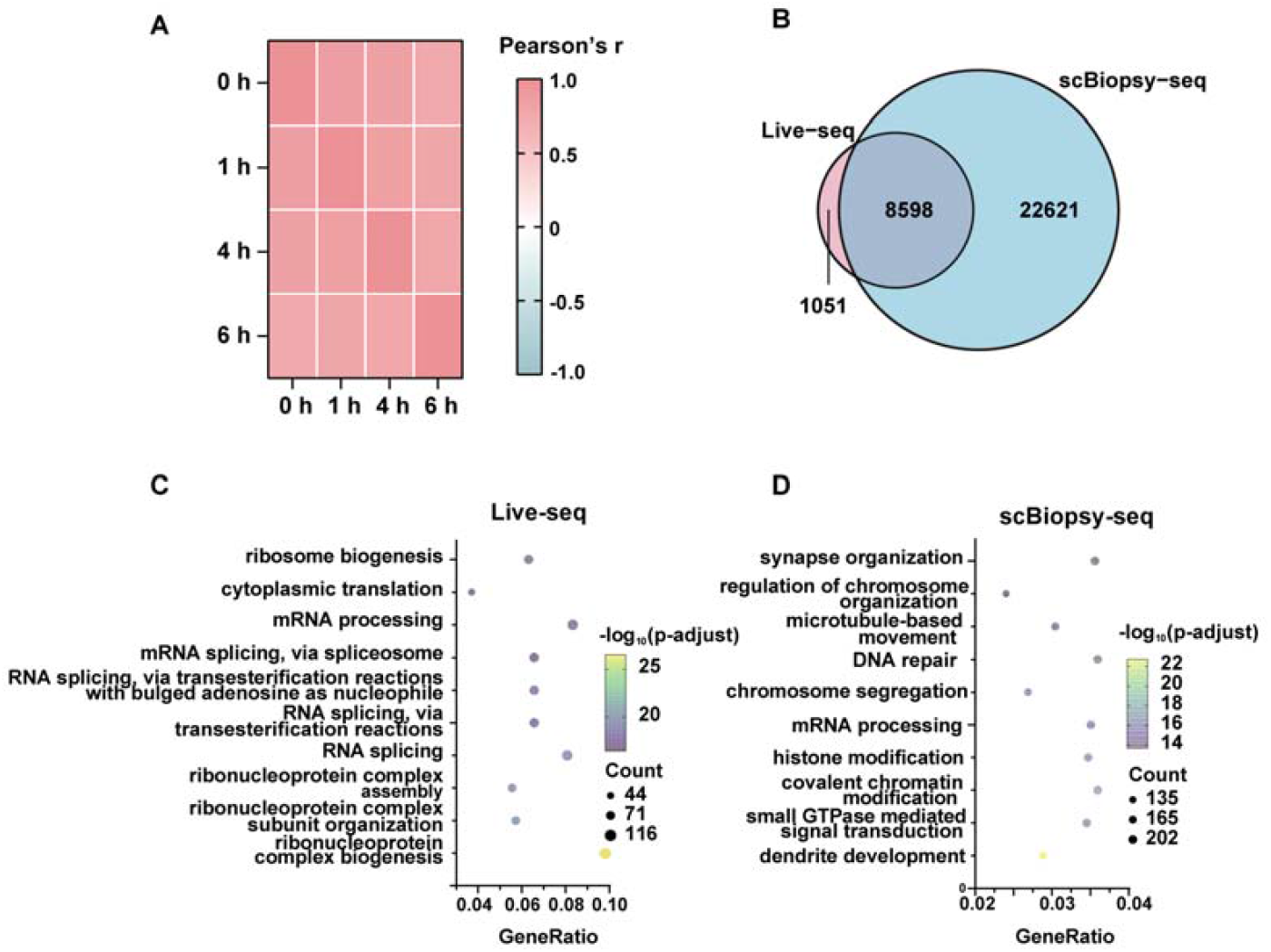
The functional reproducibility and diversity of scBiopsy-seq. **A**. Gene expression correlation (Pearson’s r) of transcriptome data from different extractions at various time points. N=7. **B**. Venn diagram exhibiting genes of RAW264.7 cells detected by Live-seq and scBiopsy-seq. All the samples were downsampled to the depth of 1 million reads. N = 10 for Live-seq and N=7 for scBiopsy-seq. **C**. Functional enrichments of genes detected by at least 50% of the extractions in Live-seq. **D**. Functional enrichments of genes detected by at least 50% of the extractions in scBiopsy-seq.

In addition, the extraction operation had no dramatic effects on cell viability (**Supplementary Fig. 4D**), number of detected genes (**Supplementary Fig. 4E**), mapping efficiency (**Supplementary Fig. 4F**) and results of clustering (**Supplementary Fig. 4G**).

We finally benchmarked the functional diversity as the potential benefits from high gene detection efficacy. Live-seq can detect 9K genes from 10 extractions. In contrast, >31K genes were detected from 7 extractions by scBiopsy-seq with significant overlap of the detected genes by Live-seq (**Fig. 4B**). The expression of the extra genes detected by scBiopsy-seq covered a broad range (**Supplementary Fig. 4H**), indicating a general improvement of data quality for scBiopsy-seq. We then interpreted the potential function of genes detected by at least 50% of the extractions with GO term enrichment. Many of the detected genes by Live-seq were associated either mRNA processing or translation, accounted for 22% of the 1450 genes (**Fig. 4C**). In contrast, 21% of the 5621 genes by scBiopsy-seq were associated with various biological processes including synapse organization, chromatin modification, histone modification, DNA repair, small GTPase mediated signal transduction, etc (**Fig. 4D**). When investigating the coverage in NF-κB signaling pathway, scBiopsy-seq can detect associated 84/104 genes (80%) while Live-seq can detect 56/104 genes (53%) (**Supplementary Fig. 4I**). All of the results suggested that scBiopsy-seq assay can generate transcriptome data with comparable quality as scRNA-seq assay in single-extraction resolution.

### scBiopsy-seq analysis identifies the mechanism for single-cell diversity in response to transcriptional suppression

The mechanism for the regulation of global transcription level is a fundamental biological question. BRD4 is a master regulator for transcription, whose inhibition induces the global suppression of transcription ^23^. In order to understand how the single-cell diversity contribute to the response to transcriptional supression, a temporal scRNA-seq analysis recording the changes of transcriptome in a time course is desired. We applied scBiopsy-seq to study this biological process with a PROTAC based chemical degrader ARV771 for BET protein ^24^. ARV771 treatment led to almost a total loss of BRD4 protein within a few hours in a VHL-proteasome dependent manner, which makes it suitable to be used for temporal scRNA-seq assay. The cell viability of a collection of on gastrointestinal cancer cell lines was used as the marker for global level of transcription in our study, since the suppression of transcription can robustly decrease cancer cell viability ^25^. We observed that the SW480 cells presented the highest IC50 and most modest response to ARV771 as well as JQ1, the original BRD4 inhibitor ^26^ (**Fig. 5A and Supplementary Fig. 5A**), indicating the potential high single-cell diversity with this cell line. We therefore investigated the temporal response of SW480 cells to ARV771 treatment with single-cell resolution by profiling the transcriptome expression at 0h, 1h and 4h, to investigate how the transcriptional dynamics influences the subsequent phenotypic responses (**Figure 5B and Supplementary Table 8**).

**Fig. 5.**
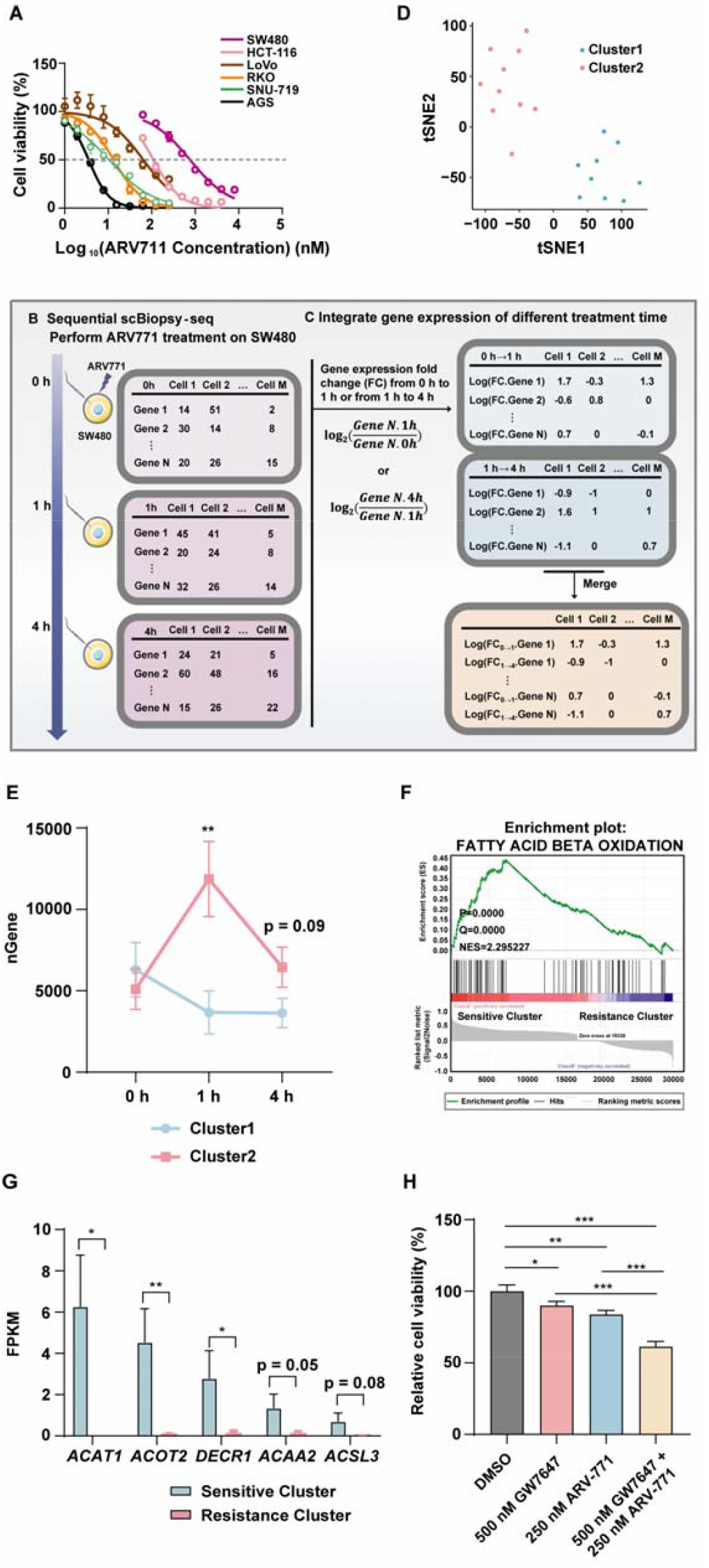
scBiopsy-seq analysis identified the mechanism of transcriptional suppression. **A**. The cell viability of different gastrointestinal cancer cells after ARV771 treatment. The IC50 values were calculated as follows: SW480: 759.30 nM; HCT-116: 110.00 nM; LoVo: 10.41 nM; RKO: 13.30 nM; SNU-719: 64.18nM; AGS: 3.55 nM. **B**. Schematic of the experimental design for sequential scBiopsy-seq analysis. **C**. Schematic of the computational algorithm for the data sequential scBiopsy-seq analysis. The differentially expressed genes in the 0-1 h and 1-4 h of the same cell were merged to generate an integrated gene expression matrix. **D**. tSNE map of integrated gene expression matrix from ARV771-treated SW480 cells by unsupervised clustering. **E**. The number of detected genes from the cells in cluster 1 or cluster 2 at 0 h, 1 h and 4 h. N=9-10. **F**. GSEA revealing significant enrichment of upregulated genes associated with fatty acid beta oxidation in sensitive cluster compared to resistance cluster. **G**. The expression of genes encoding key factors for fatty acid beta oxidation in the cells of sensitive cluster or resistance cluster. N=9-10. **H**. Relative cell viability of SW480 cells after the treatment of ARV771, PPARa agonist GW7647 or in combination. N=5.

In order to analyze the high-dimensional data in a time course, we designed a computational algorithm for the multi-modular dataset, which integrated the differentially expressed genes in the 0-1h and 1-4h matrices of the same cell (**Figure 5C and Supplementary Methods**). The changes of transcriptome in the time course were clustered into two groups (**Figure 5D**). The number of detected genes was downregulated in cluster1 but not cluster2 by ARV771 treatment for 1h and 4h (**Figure 5E**). In consistent to the existing knowledge that BRD4 is an important regulator for the transcriptional activator MYC ^27^, the functional enrichment revealed that a large part of significantly downregulated genes between 1h and 0h in cluster1 were MYC-targeted (**Supplementary Fig. 5B-C and Supplementary Table 9**). These results indicated that the cells in cluster1 are more sensitive to BRD4 degradation induced transcriptional suppression.

Based on the theory of clonal evolution, the temporal response to transcriptional suppression in single-cell resolution is decided by the biological features of cells before treatment. The gene set enrichment analysis (GSEA) on the differentially expressed genes at 0h between sensitive cluster and resistance cluster discovered that the enrichment of genes related to fatty acid beta oxidation in the sensitive cluster (**Fig. 5F**). The expression of multiple genes, encoding the key enzymes for fatty acid beta oxidation, was downregulated in the cells of resistance cluster (**Fig. 5G**). The combinational treatment of ARV771 and PPARa agonist GW7647 ^28^, which is a widely accepted enhancer for fatty acid beta oxidation ^29^, upregulated the expression of related genes (**Supplementary Fig. 5D-E**) and prevented the growth of SW480 cells (**Fig. 5H**). These results indicated that the application of scBiopsy-seq assay identified an unconventional contribution of lipid metabolism to the BRD4 degradation induced transcriptional suppression.

## Discussion

In the past decades, many efforts have been made to obtain more information from single-cell technology. As all these assays need to destroy the cells in order to get the information, it is a great challenge to capture the features of one cell for multiple times. In the current study, we established an assay called scBiopsy-seq to measure the transcriptome of single cell in a time course, which can perform the sequential extraction of the cytoplasm from the same cell multiple times, enabling the association of the cell phenotypic responses with its transcriptional dynamics. In comparison to the existing temporal scRNA-seq assay, scBiopsy-seq can extract the samples with well-controlled volume by synergistic EOF-EP, which improves the reliability of the experiment. Furthermore, the use of oil-isolated hydrophobic interface on DMF for RNA library processing enhances the detection sensitivity and accuracy. The enhanced detection efficacy of scBiopsy-seq, on both the number and functional diversity of detected genes, demonstrated the dramatic improvement of data quality. The employment of scBiopsy-seq on the study of transcriptional regulation revealed a lipid metabolism related mechanism for the resistance to BRD4 inhibition.

The establishment of scBiopsy-seq assay may expand the application of temporal scRNA-seq in the laboratories focusing on solving cell biological questions. As we designed it as a flexible platform, biologists can build their own scBiopsy-seq platform with regular equipment. All the reagents or consumables for scBiopsy-seq assay can be either obtained from commercial vendors or generated with regular equipment based on the experimental details we provided. With scBiopsy-seq assay, it is possible to understand more complicated biological processes, such as the responses to hormones/nutrients stimulation in primary hepatocytes or liver organoids. Although we only validated the efficacy of scBiopsy-seq with mammalian cells, its application in the field of microbiology (such as the response of yeast to environmental challenges ^30^) or plant biology (such as the response to light stimulations ^31^) is expected.

In summary, scBiopsy-seq assay can perform the extraction and measure the transcriptome of single cell reliably and efficiently. The quality of the transcriptome data from scBiopsy-seq is comparable to SS2 based scRNA-seq assay. The establishment of scBiopsy-seq assay expands the application of temporal scRNA-seq assay, which may bring the community of single-cell technology to enter the age of real-time.

## Declaration of interests

None

## Author contributions

Conceptualization, L.C., S.L., M.Q., L.L., F.L., J.L., Y.Z., X.N., S.L., X.X., C.Y. and J.L.; Investigation, L.C., S.L., M.Q., L.L., F.L., J.L., Y.Z., X.N., S.L., X.X., C.Y. and J.L.; Analysis, L.C., S.L., M.Q., L.L., F.L., J.L., Y.Z., X.N., S.L., X.X., C.Y. and J.L.; Writing, L.C., S.L., M.Q., L.L., F.L., J.L., Y.Z., X.N., S.L., X.X., C.Y. and J.L.; Data Visualization, L.C., S.L., M.Q., L.L., F.L., J.L., Y.Z., X.N., S.L., X.X., C.Y.x and J.L.; Funding Acquisition, X.X., C.Y. and J.L.; Supervision, X.X., C.Y. and J.L..

## Acknowledgments

This work was supported by MOST 2023YFA1801000, MOST 2020YFA0803601, NSFC 32371193, NSFC 32071138 and STCSM 23XD1420300 to Jin Li, NSFC 22293031 to Chaoyong Yang and NSFC 22204132 to Xing Xu.

## Data availability

Sequence data that support the findings of this study have been deposited in in the GEO database with the accession number GSE283737 (please use the token ixspgqymzxyjpan). The code used in this study is available on https://github.com/linl-linl/scBiopsy-seq.

## Notes

### Competing Interest Statement

The authors have declared no competing interest.

