## Supplementary File for "scBiopsy-seq: a platform for temporal single-cell RNA-seq analysis"

### Supplementary Materials and Methods

#### Cell culture

The RAW264.7, SW480, HeLa and SK-BR-3 cells were purchased from National Infrastructure of Cell Line Resource and cultured in an incubator with 5% $CO_2$ at 37°C. RAW264.7 and SW480 cells were cultured in high-glucose DMEM medium containing 10% fetal bovine serum (ThermoFisher Scientific Co., Ltd., Shanghai, China) and 1% penicillin-streptomycin solution (ThermoFisher Scientific Co., Ltd., Shanghai, China). HeLa cells were cultured in MEM medium containing 10% fetal bovine serum and 1% penicillin-streptomycin solution. SK-BR-3 cells were cultured in McCoy's 5A medium containing 10% fetal bovine serum and 1% penicillin-streptomycin solution.

#### Fabrication of digital microfluidic chip

The DMF device comprised a top plate (75 mm × 30 mm × 1.1 mm) and a bottom plate (80 mm × 63 mm × 1 mm). The top plate was based on an indium tin oxide (ITO) glass, which was initially ultrasonically washed with glass cleaning solution at 65°C 45 minutes, rinsed in water, and dried for utilization. Subsequently, the top plate was coated with a 1% (w/w) Teflon-AF solution by spinning at 1500 rpm for 60 seconds, followed by heating at 200°C for 20 minutes. This operation yielded a clean top plate devoid of any patterns. The bottom plate was grounded on a glass base and had the electrode patterns designed thereon. The cleaning procedures of the substrate glass were identical to those of the top plate. Then the substrate glass was coated with a 300-nm chromium layer and underwent photolithography to form an electrode layer. Afterward, SU8-2015 photoresist was spin-coated onto the electrode layer to form a dielectric layer for alleviating the breakdown of the electrode. Finally, 1% (w/w) Teflon-AF was spin-coated onto the dielectric layer, generating a hydrophobic layer that could reduce nucleic acid adsorption. The top plate and the bottom plate were stored at room temperature.

Before utilization, both plates were rinsed with ethanol and exposed to ultraviolet light. Subsequently, the two plates were aligned with a spacer (120 μm) in between, which was formed by applying two layers of single-sided tape to each side of the bottom plate. During the processing of the RNA library, this gap was filled with PMX-200 silicone oil (S104740-1L, Aladdin Biochemical Technology Co., Ltd., Shanghai, China) to create a fully sealed environment. The chip was placed on a chip holder equipped with 48 pogo pins, which connected the DMF chip and the home-made control box. The control box output electrical signals to charge or discharge electrodes, enabling the precise manipulation of droplets.

**Quantification method of extraction volume**

To determine the order of magnitude range of the extraction volume, we introduced the concept of electrochemical equivalent volume (EEV). Electrochemical equivalent volume refers to the volume represented by the amount of electricity generated during the extraction of potassium ferrocyanide standard electrode pairs in a cell environment. Specifically, after a given extraction voltage and time, the potassium ferrocyanide standard electrode pair will generate a certain amount of electricity, and the corresponding volume can be calculated by Faraday's law:

$$EEV = Q/nCF$$

where Q is the amount of electricity measured by the electrochemical method, n is the number of charges transferred during the process (for potassium ferrocyanide standard electrode pairs, n = 1), C is the molar concentration of potassium ferrocyanide, and F is the Faraday constant (96485 C/mol).

The EEV measurement was performed using chronocoulometry with CHI1200C (Shanghai, China). Specifically, the standard electrochemical solution (1 M potassium ferricyanide/potassium ferricyanide solution (Aladdin Biochemical Technology Co., Ltd., Shanghai, China) and 0.1 M KCl (Beyotime Biotechnology Co., Ltd., Shanghai, China) as the supporting electrolyte) was employed as redox pairs. The 0.1 M KCl solution was filled in the capillary and served as the work electrode. The chronocoulometry was performed at different times (3, 5, 10, 15, 20, 30, 40, 60, and 80 seconds) in accordance with the three-electrode connection and different voltages (0.3, 0.5, 1.0, 2.0, and 3.0V *versus*. Ag/AgCl).

It is notable that due to the presence of bioelectrical impedance in actual samples and the difference in viscosity and mobility between the extracted material and potassium ferrocyanide, the actual extraction volume is typically smaller than EEV. Nevertheless, it meets the requirements to evaluate the damage to cells during the extraction process when the EEV is higher than the actual volume.

**Preparation of APTES-modified glass capillary**

The nanopipettes (length l = 8.50 cm, outer diameter o.d. = 1 mm, inner diameter i.d. = 0.7 mm, Huaicheng Quartz and Micrology precision Instruments Co., Ltd., Wuhan, China) with ~300 nm needle tips were fabricated by the laser pulling instrument (P2000; Sutter Instrument Co., Ltd., Novato California, America). The pulling parameters were: Line 1: Heat 850, Fil 4, Vel 12, Del 128, Pul 140; Line 2: Heat 850, Fil 4, Vel 15, Del 132, Pul 210. The needle tip was characterized by scanning electron microscope (SEM). For the covalent modification of APTES (91930-2, Aladdin Biochemical Technology Co., Ltd., Shanghai, China) on the surface, the nanopipettes were then acidified to expose silanol groups on the surface. This procedure encompassed immersing the nanopipettes in a diluted nitric acid solution ($HNO_3$, 6.5%, 40 mL), and subsequently subjecting them to sonication using an ultrasonic bath at 70°C for 10 minutes. After acidification, the nanopipettes were rinsed three times with ultra-pure water (40 mL) before being dried at 100 °C for 2 hours. Subsequently, the acidified nanopipettes were immersed in APTES (99%, 40 mL) and subjected to sonication at 70 °C for 3 hours. Upon completion of the sonication, the nanopipettes were washed three times with ethanol (95%, 40 mL) and ultra-pure water (40 mL), and finally dried at 100 °C for 2 hours, yielding amino-carried glass capillary.

**The simulation for fluid physics**

The establishment of the numerical model involved several key steps aimed at understanding the behavior of RNA molecules within a capillary under specific conditions. The process included defining the geometric structure, selecting appropriate control equations, setting boundary conditions, and performing simulations to analyze the outcomes.

The initial step was to define the geometry of the system, which comprised a conical nanocapillary (ABCF) with a small fluid reservoir outside the capillary (FCDE) and an Ag/AgCl electrode (AGHB). The dimensions were specified as follows: AB = 8 μm, AF = BC = 30 μm, and EF = FC = CD = ED = 300 nm.

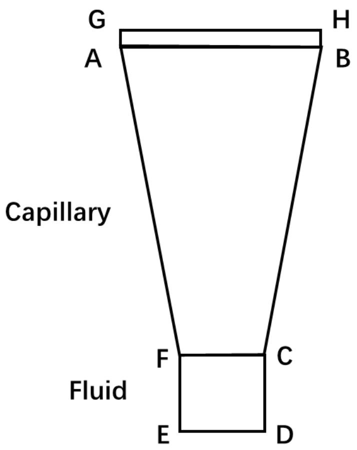


Boundary conditions were established to accurately reflect the operational environment. These included: (a) On AB, a voltage of -0.5 V or +0.5 V was applied; (b) On ED, zero charge condition was assumed; (c) AF & BC had a surface charge density of -1 $mC/m^2$; (d) EF & CD had zero charge; (e) At points F and C, the voltage was set to 0 V; (f) For all boundaries, the RNA concentration was initially 1 mM, and the concentrations of $Na^+$ and $Cl^-$ were equal; (g) No-slip conditions were enforced at the solid interfaces.

The electrochemical trapping process was numerically simulated using COMSOL Multiphysics 5.3a. A 2-D axisymmetric model was developed, employing three modules: "Transport of Diluted Species", "Electrostatics", and "Laminar Flow". The Nernst-Planck, Poisson, and Navier-Stokes equations [1] were utilized to simulate mass transport, electric field distribution, and fluid dynamics within the nanocapillary, respectively. Boundary conditions were defined as follows:

| Boundary | Poisson | Nernst-Planck | Navier-Stokes |
|---|---|---|---|
| AB | V=-0.5\0.5V | c(RNA)=1mM<br>$c(Na^+)=c(Cl^-)=1mM$ | (−∇ + ∇ 2 ) · = 0 |
| ED | Zero charge | c(RNA)=1mM<br>$c(Na^+)=c(Cl^-)=1mM$ | (−∇ + ∇ 2 ) · = 0 |
| AF&BC | -1 mC/m2 | No flux | = 0 |
| EF&CD | Zero charge | No flux | = 0 |
| Point F&C | V = 0 V | No flux | = 0 |

Simulations were conducted under varying voltages and surface charge densities. Key parameters analyzed included the concentration profile of RNA within the capillary, the velocity field, and the volumetric flow rate. These analysis revealed that changes in voltage and surface charge significantly impacted the RNA concentration distribution and fluid dynamics within the capillary.

**The process of scBiopsy-seq**

The capillary tip was filled with 1.5 μL of mixed solution (2.5 μM biotinylated-oligo-dT primer, 2.5 mM dNTP (AD101–12, TransGen Biotech, Beijing, China), 0.4 $U \cdot \mu L^{-1}$ Murine RNase Inhibitor (GMP4102PA-01, Vazyme Biotech Co., Ltd., Nanjing, China)), and then connected to the work electrode of the ultra-micro extraction instrument (Rayme Biotechnology Co., Ltd., Jiangsu, China). The counter electrode and the reference electrode were immersed in the culture medium within the confocal dish. The cells to be extracted were visualized by microscope and the preloaded capillary was gradually moved downward until it touched the cytoplasm of the selected cell. The tip of the capillary was then inserted into the cytoplasm, with a deformation observed in the selected cell. By setting the applied voltages and the duration of current application, a precisely controlled volume was extracted from the cell. After that, the capillary was retracted out of the cell, taken out of the extraction instrument and placed at -80°C for RNA storage.

Once it was required to undergo the RNA library construction, the capillary was connected to the work electrode again. The cellular contents within the capillary were injected into the top plate of the DMF device, enabling the subsequent RNA library processing on the DMF system. The lysis buffer (2.5 μM biotinylated-oligo-dT primer, 2.5 mM dNTP, 0.4 U·μL$^{-1}$ Murine RNase Inhibitor and 0.2% Triton-X-100(A110694, Sangon Biotech, Shanghai, China) was introduced and mixed with the cellular contents at 72°C for 3 minutes for complete RNA release. Then the reverse transcription (RT) mix (1.7 μM biotinylated-TSO oligo, 5 × HiScript II RT buffer (Vazyme Biotech Co., Ltd., Nanjing, China), 1.7 U·μL$^{-1}$ Murine RNase Inhibitor, 19 U·μL$^{-1}$ HiScript II Reverse Transcriptase (R201, Vazyme Biotech Co., Ltd., Nanjing, China) and 0.05% Pluronic F68 (24040032, ThermoFisher, America)) was introduced to initiate the RT reaction at 42°C for 90 minutes, which was stopped by heat at 70°C for 15 min. After that, the PCR mix (2 × KAPA HiFi HotStart ReadyMix (KK2602, KAPA, America), 0.06 μM Biotinylated-ISPCR oligo and 0.05% Pluronic F68) was introduced to amplify the cDNAs (95°C 3 minutes, 25 cycles of 98°C 30 s, 65°C 45 s and 72°C 6 minutes, followed by a final step at 72˚C for 5 min). The products were recovered from the DMF device into a 200-μL eppendorf tube, purified with 0.6 × DNA clean beads (Vazyme Biotech Co., Ltd., Nanjing, China) and quantified by qubit 3.0. Then 1 ng of the products were utilized and processed by sequencing-ready library construction kits (TD503, Vazyme). The library was purified twice with 0.6 × DNA clean beads (N411, Vazyme Biotech Co., Ltd., Nanjing, China) and sequenced by Novaseq500 according to the manufacturer's instructions.

For sequential scBiopsy-seq, cellular contents of the same cell was extracted at different time points. All of the steps were monitored in real time by microscopy in bright field.

**The process of scRNA-seq**

Cells were digested with trypsin and dissociated into a single-cell suspension, and subsequent processing followed the same steps as DMF-RNA-seq [2] which implemented Smart-seq2 on the DMF chip. Specifically, single cells were captured by hydrophilic site on a DMF chip, and then lysed to completely release RNAs. By introducing the lysis buffer, RT mix and PCR mix in sequence, amplified cDNAs were obtained. The cDNAs were then purified, processed using sequencing-ready library construction kits (TD503, Vazyme Biotech Co., Ltd., Nanjing, China) and eventually sequenced by Novaseq500.

**The treatment of BRD4 inhibition on cells**

The BRD4 inhibition ARV771 was purchased from TargetMol (Cas1949837-12-0), and JQ1 was purchased from MCE (HY-13030). To study the temporal response of SW480 cells to ARV771 treatment, sequential scBiopsy-seq was processed on single SW480 cells. The cellular contents of SW480 cells were extracted initially prior to the ARV771 treatment, designated as the sample at 0 h. Immediately thereafter 250 nM ARV771 was added to the culture medium, and the cells were extracted at 1 h and 4 h.

**Bulk RNA-seq assay**

Total cells RNA was extracted using the Zymo Quick-RNA miniprep kit (Zymo, #R1055) according to the manual. PolyA mRNA was purified using NEBNext PolyA mRNA Isolation Module (NEB, #E3370S), as per the manufacturer's instructions. RNA-seq libraries were prepared using the NEBNext Ultra Directional II RNA library prep kit and sequenced by Illumina/BGI with a 150 bp paired-end run configuration. Three biological replicates per sample.

Quality control was performed using the FastQC (v0.11.7) tool and the results were analyzed. Cleaned RNA-seq reads were mapped using STAR[3] (v. 2.7.10) against the human genome (hg19). Uniquely mapped reads were quantified with HTSeq[4] (v.0.11.3) and protein-coding genes with nonzero read count (n = 20138) were included for downstream analysis. Differential expression of protein-coding genes was analyzed with these counts using Bioconductor package DESeq2[5] (v.1.40.1). Significantly differential expression was considered by setting adjP < 0.05 and FC ≥ 1.5 between sh*MYB* and shNC. Gene Ontology (GO) and KEGG analysis were performed using the online DAVID tool[6]. The gene set enrichment analysis was performed according to the instructions (https://www.gsea-msigdb.org/gsea/index.jsp).

**Cell growth assay**

For cell growth assays, selected cells were collected. 5000 cells were seeded into 96-well plates (100 μL/well). After 5 days, 10 μL of Cell Counting Kit-8 (CCK-8) (Beyotime, #C0040) was added into each well according to the manufacturer's instructions and incubated for 1 h. Optical density was measured at 450 nm using a microplate reader.

**Sequencing data quality control**

Cutadapt v3.4 was employed to remove bases and adapter sequences with a sequencing quality lower than 20 at the 3' end of the sequence, and sequences containing 30 identical polyA/G/T bases were discarded. Subsequently, repair.sh in BBmap v38.90 was utilized to repair the sheared sequence.

GRCh38 and GRCh38.105 were utilized as the reference genome and reference genome annotation files for human samples. GRCm38 and GRCm38.102 were employed as the reference genome and reference genome annotation files for mouse samples. STAR v2.7.3a was used to map the repaired sequence to the reference genome. The file output by the software could obtain information such as the unique alignment rate and the multi-site alignment rate.

Exon/intron alignment rate: Information entries with feature types of "gene" and "exon" in the reference genome annotation file were extracted and organized into bed files containing the feature types of each site. Samtools v1.3.1 was employed to refer to the information provided by the bed file to calculate the sequencing depth of each site on the whole genome, exon region, and gene fragment. The sum of the sequencing depths of all sites on the feature type was regarded as the number of bases detected for the feature type. The number of bases in the intron region was obtained by subtracting the number of bases in the exon region from the number of bases on the gene fragment. Finally, the ratio of the number of bases of each feature type to the number of bases in the whole genome was computed as the exon alignment rate, intron alignment rate and gene alignment rate.

Number of genes: The alignment result sam file was filtered to the same depth (1 million Reads) by random sampling, and the number of reads of each gene in the alignment was counted using htseq-count v0.12.4. The number of genes with non-0 reads was the number of genes detected at this depth.

Mitochondrial expression (proportion): The gene name/gene id of mitochondrial-related genes was acquired from the reference genome annotation file. The total expression/reads of these genes constituted the mitochondrial expression, and the ratio of the total number of mitochondrial reads to the total number of cell reads was the mitochondrial expression ratio.

**Cell clustering**

All samples were down-sampled to the same depth, and the gene expression of each sample was calculated by htseq-count v0.12.4 and integrated into an expression matrix. The R package SC3 v1.18.0 was used to filter the expression matrix according to the default percentage, and the cell clustering analysis was performed using the k-means algorithm. The clustering results were presented in the form of t-SNE dimensionality reduction diagram or heat map by the plotTSNE function or sc3_plot_expression function of SC3. SC3 clustering parameters: biology=TRUE, gene_filter=T, d_region_min=0.06, d_region_max=0.25, pct_dropout_min=20, pct_dropout_max=80.

**Pearson correlation analysis**

The samples on different platforms or at different time points were preprocessed as above to obtain the gene expression matrix, and the mean of the gene expression was taken as the expression detected on each platform/time point. In order to avoid the influence of high expression data, the expression count value was logarithmized. The Pearson correlation coefficient was used to evaluate the similarity of expression levels detected on different platforms/time points for all genes.

**Heat map and GO enrichment analysis based on gene expression at different time points**

The analysis process of the differential expression referred to that in live-seq [7]. Specifically, the gene expression of the samples to be compared was extracted, and gene filtering was first performed to retain genes with an expression level $\geq 2$ in $\geq 5\%$ of the cells and genes with an expression level $\geq 2$ in more than 15% of the samples at any time point. Then, the R package edgeR (v3.34.1) was used for differential expression analysis. Once the library was standardized, the dispersion was estimated and the negative binomial distribution (NB) model was fitted using the quasi-likelihood F-test, enabling to obtain the differential expression significance p value and the logarithm of the differential expression multiple logFC for each gene. The significant p value was then corrected using the Benjamini-Hochberg method. In order to ensure the effectiveness of the differential analysis, genes whose expression level was greater than 15% in both the target time point sample and other samples were filtered out. Subsequently, $p < 0.05$, $|logFC| > 1$ were used as thresholds to identify differentially expressed genes, where $logFC > 0$ was upregulated genes, and vice versa, downregulated genes. Based on differentially expressed genes, R package clusterProfiler v3.18.1 was used for functional enrichment analysis. With the GO database as a reference, upregulated/downregulated genes were used as analysis objects to enrich biological function modules that changed with drug administration time. Parameter was set as: fun="enrichGO", OrgDb='org.Hs.eg.db', pvalueCutoff=0.05.

**The integration of the matrix of differential gene expression changes at different time points**

First, the expression level of each cell at 0 h was used as the benchmark, and the expression level of 1 h was normalized by calculating the expression change fold of 1 h relative to 0 h. Similarly, the expression change fold of 4 h relative to 1 h was calculated. The expression changes in different time periods were regarded as independent cell features, and the feature matrix of cell, behavior time period-gene expression changes was generated by integration, and the features that changed in at least 90% of the cells were retained. The filtered feature matrix was conducted principal component analysis (PCA), and the first two principal components (PCs) containing the most variation information were selected for k-means clustering (k=2). Based on the obtained cell clustering results, the Wilcoxon test was performed between the features of different cell clusters to find gene expression changes closely associated with cell clustering. Each feature (time period-gene expression change) was sorted according to the p value obtained by the test, and features with $p<0.05$ were considered to be iconic features, that is, features with significant differences between cell clusters.

**Statistical analysis**

All statistical comparisons between two groups were performed by GraphPad Prism software 9.0 using a two-tailed unpaired *t*-test, unless specified. The variance between the statistically compared groups was similar.

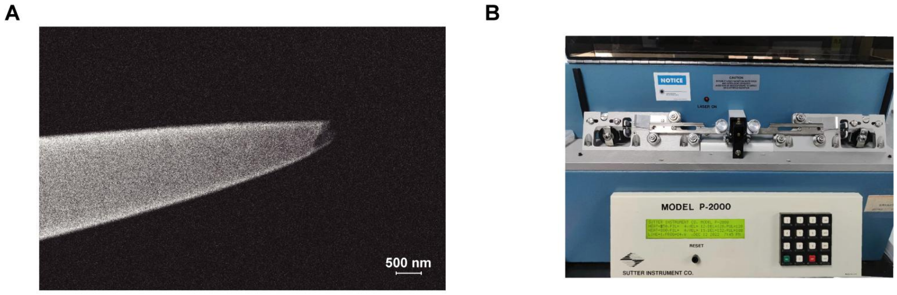


**Supplementary Fig. 1 The instruments to perform extractions for scBiopsy-seq.**

**A.** The scanning electron microscope image of the glass capillary with a 300 nm needle tip used for extraction.

**B.** The instrument and parameter setting for creating the glass capillary with a 300 nm needle tip.

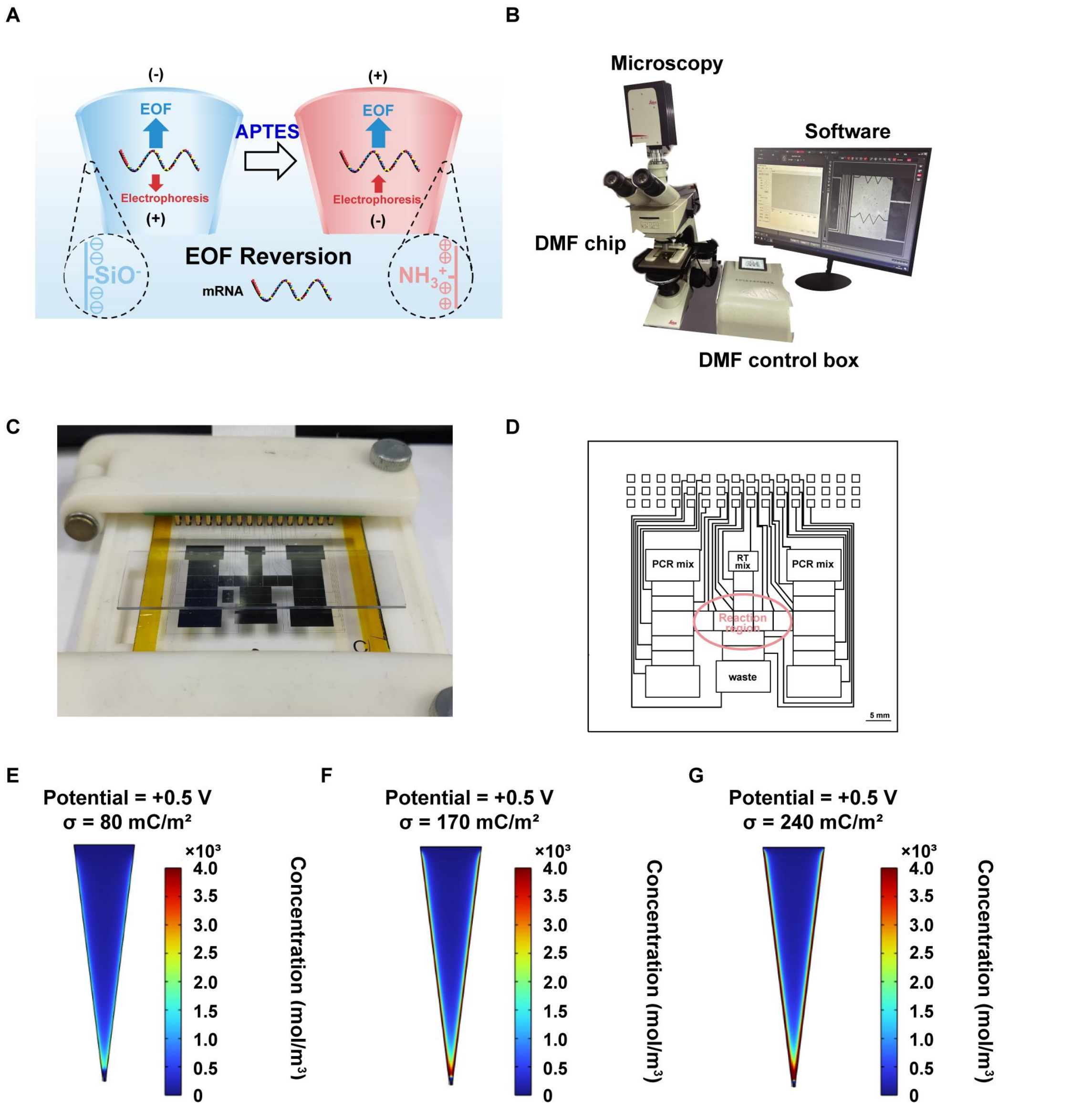


**Supplementary Fig. 2 The synergistic electroosmosis and electrophoretic force and microfluidics instruments for scBiopsy-seq.**

**A.** The principle for the synergistic effects of EOF and EP by APTES modification.

**B.** The integrated digital microfluidics system with microscope, digital microfluidic chip, DMF control box and microfluidic software, which was constructed for automatic RNA sample processing.

**C.** The physical DMF chip fixed in a chip holder.

**D**. The computer aided design of chip design for scBiopsy-seq, including the reservoirs for the introduction of reverse transcription mix and polymerase chain reaction mix, as well as the reaction region for RNA library processing.

**E.** Fluid physics simulation of concentration field with COMSOL Multiphysics within a nanopipette during the extraction process. The charge surface density ($\sigma$) was 80 $mC/m^2$ and the corresponding applied potential was +0.5 V. The extracting concentration of negatively charged molecules, such as RNAs, was presented in color scale. Average concentration of negatively charged substance within the nanopipette was calculated at 367.52 mM.

**F.** Fluid physics simulation of concentration field with COMSOL Multiphysics within a nanopipette during the extraction process. The $\sigma$ was 170 $mC/m^2$ and the corresponding applied potential was +0.5 V. The extracting concentration of negatively charged molecules, such as RNAs, was presented in color scale. Average concentration of negatively charged substance within the nanopipette was calculated at 834.25 mM.

**G.** Fluid physics simulation of concentration field with COMSOL Multiphysics within a nanopipette during the extraction process. The $\sigma$ was 240 $mC/m^2$ and the corresponding applied potential was +0.5 V. The extracting concentration of negatively charged molecules, such as RNAs, was presented in color scale. Average concentration of negatively charged substance within the nanopipette was calculated at 1215.40 mM.

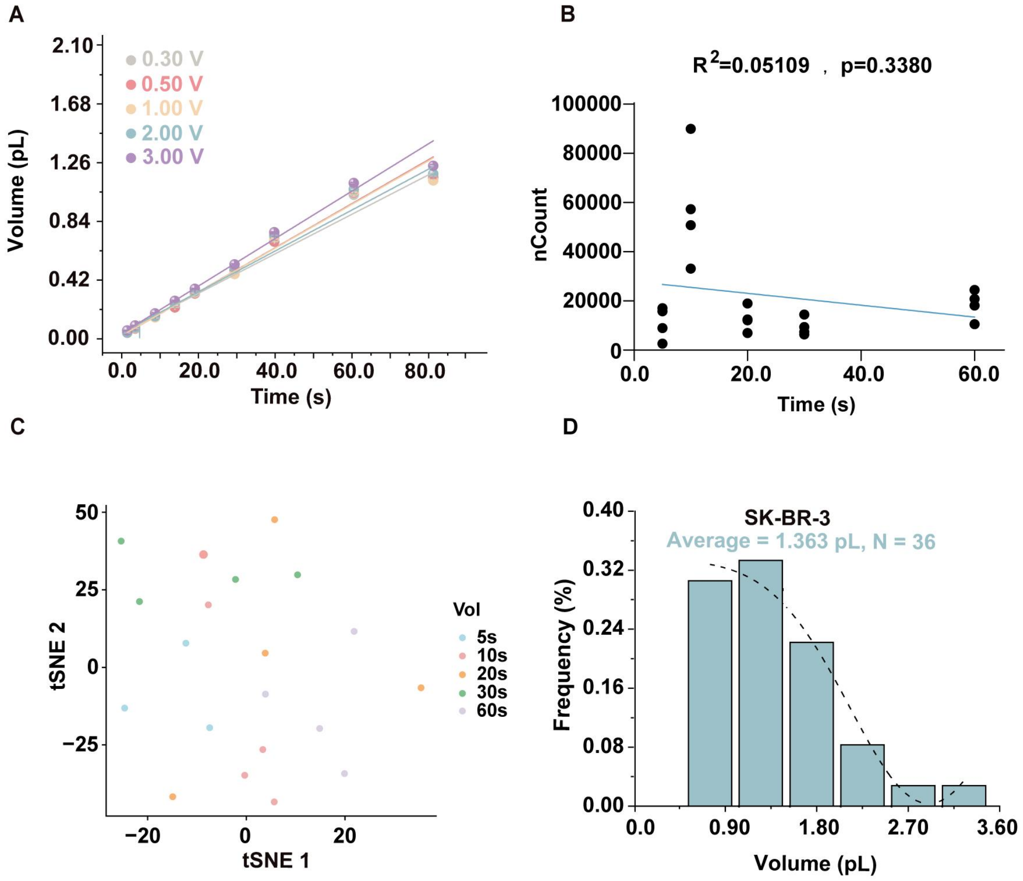


**Supplementary Fig. 3 The well-controlled extraction volume of scBiopsy-seq.**

**A.** Calibration curve of extraction volume based on extraction time. The applied potential for calibration curve varied from 0.3 V to 3.0 V (from top to bottom: y = 0.01698 x + 0.001981, $R^2$ = 0.9895 for 3.0 V; y = 0.01576 x + 0.005751, $R^2$ = 0.9887 for 0.5 V; y = 0.01565 x + 0.007381, $R^2$ = 0.9826 for 1.0 V; y = 0.01465 x + 0.002793, $R^2$ = 0.9844 for 2.0 V; y = 0.01413 x + 0.02918, $R^2$ = 0.9929 for 0.3 V, respectively). Note that the actual electrical parameters utilized in the scBiopsy-seq were set at 0.5 V for 10 seconds. N = 5.

**B.** The correlations ($R^2$ of linear regression and p value (two-sided F-test) ) between the number of counts and extraction time. N = 4.

**C.** tSNE map of transcriptome from different extraction time.

**D.** Histogram depicting the frequency distribution of cell volume of SK-BR-3 cells.

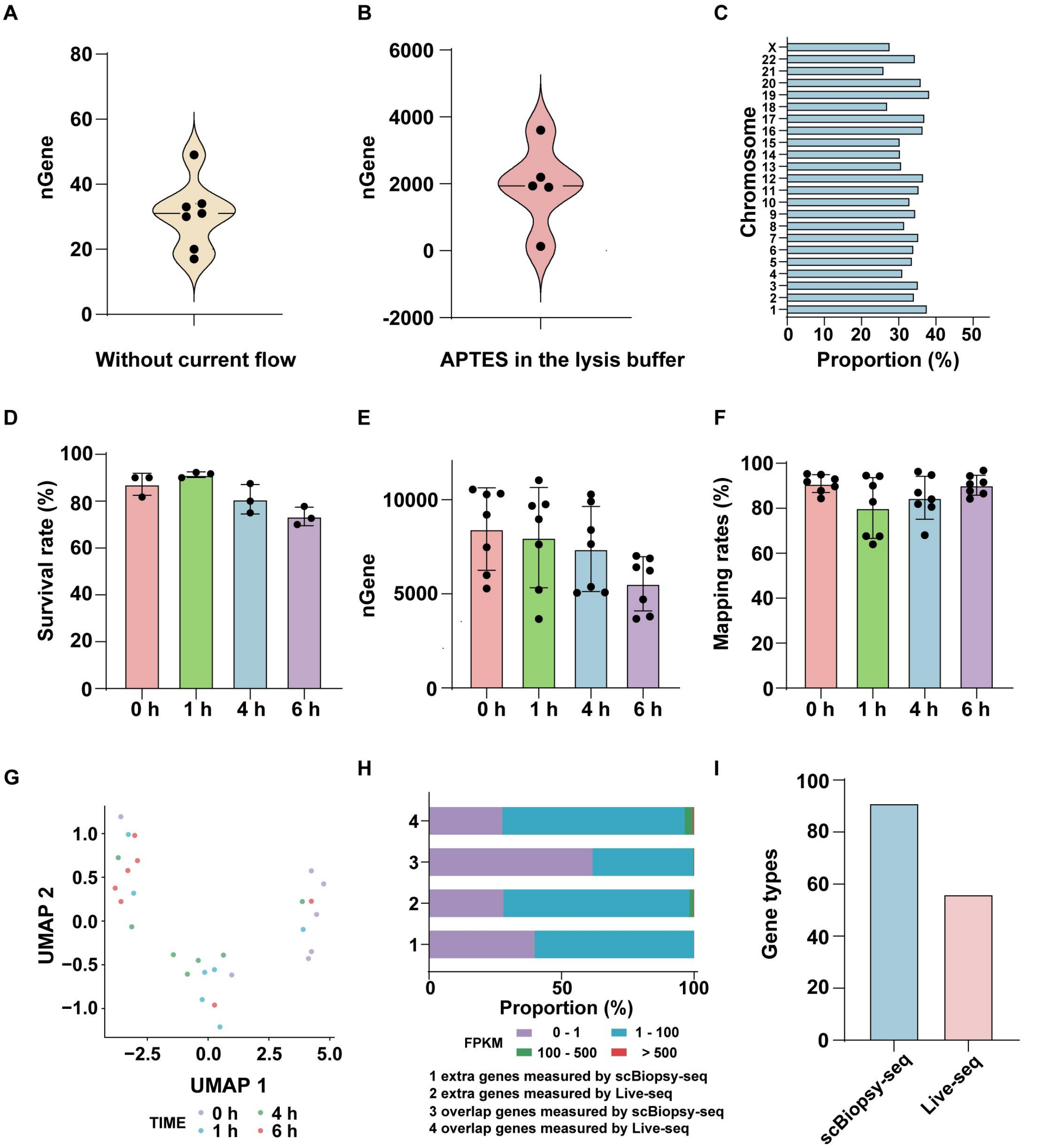


**Supplementary Fig. 4 The performance of scBiopsy-seq on measuring transcriptome.**

**A.** The number of genes detected by scBiopsy-seq in the absence of current flow. N = 7.

**B.** The number of genes detected by scBiopsy-seq when APTES was directly added to the lysis buffer instead of covalent modification on the glass capillary. N = 5.

**C.** The distribution of detected genes by scBiopsy-seq in different chromosomes.

**D.** The survival rate of SK-BR-3 cells after extraction at various time points. Three groups were tested and 20 cells were contained in each group.

**E.** The number of detected genes by scBiopsy-seq at various time points in the SK-BR-3 cells. N = 7.

**F.** The mapping efficiency of data derived from SK-BR-3 cells by scBiopsy-seq at various time points. N = 7.

**G.** tSNE map of transcriptome data derived SK-BR-3 cells by scBiopsy-seq at various time points.

**H.** The expression levels (FPKM) of genes detected by scBiopsy-seq or Live-seq, based on the analysis of **Fig. 4B**.

**I.** The gene types of NF-κB pathway in RAW264.7 cells detected by scBiopsy-seq (N = 7) or Live-seq (N = 10).

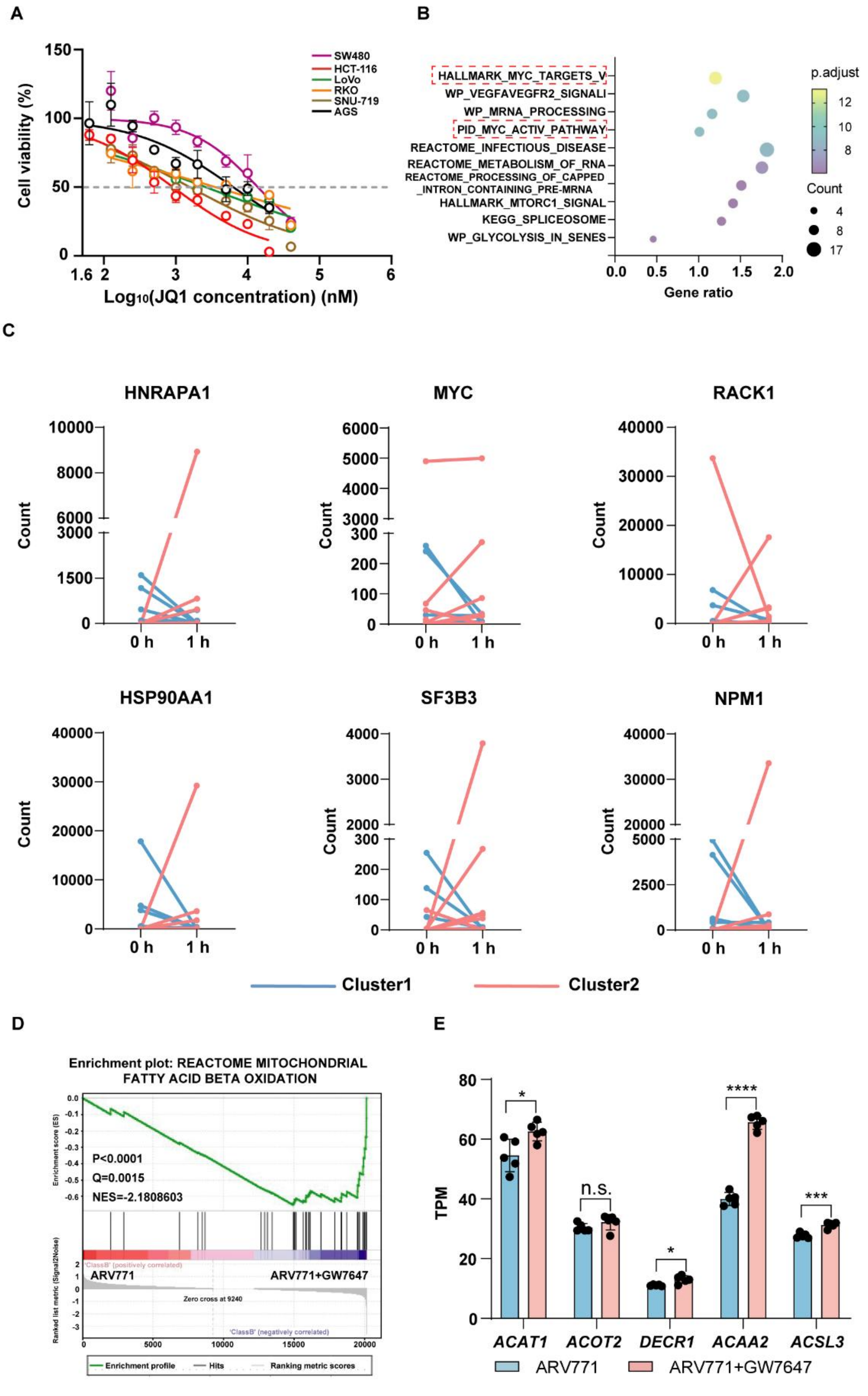


**Supplementary Fig. 5 scBiopsy-seq analysis on transcriptional suppression.**

**A.** The cell viability of different gastrointestinal cancer cells after JQ1 treatment. The IC50 values were calculated as follows. SW480: 13.833 μM; HCT-116: 0.915 μM; LoVo: 2.602 μM; RKO: 3.552 μM; SNU-719: 1.567 μM; AGS: 6.297 μM. N = 5.

**B.** Functional enrichment analysis of the significantly downregulated genes by 1-h ARV771 treatment in cluster1.

**C.** The dynamic changes of the expression for MYC targeted genes from 0 h to 1 h in cluster1 (blue) and cluster2 (red).

**D.** GSEA revealing significant enrichment of upregulated genes by the combination of ARV771 and GW7647 associated with fatty acid beta oxidation. N = 5.

**E.** The upregulation of genes related to fatty acid oxidation in SW480 cells treated with the combination of ARV771 and GW7647. N = 5.
